# Destabilization of Helix III Initiates Early Serum Amyloid A Misfolding by Exposing Its Amyloidogenic Core

**DOI:** 10.1101/2025.05.27.656333

**Authors:** Haidara Nadwa, Z. Faidon Brotzakis, Annalisa Santucci, Daniela Braconi, Michele Vendruscolo

## Abstract

Serum amyloid A (SAA) is the principal precursor of AA amyloidosis, yet the early molecular steps triggering its pathological misfolding remain unclear. Here, we combine harmonic linear discriminant analysis (HLDA) and parallel-tempering metadynamics (PT-MetaD) to dissect the earliest conformational transitions of the disease-relevant SAA_1−76_ fragment. By constructing an optimized one-dimensional collective variable (sHLDA) from inter-helix contacts and helical root-mean-square deviations, we perform 4 *µ*s of enhanced sampling across 79 replicas (300–450K). Free-energy surfaces reveal a misfolding trajectory where helix III destabilizes first, preceding loss of helices II and I while global compactness persists. Solvent-accessible surface-area analysis reveals transient exposure of the aggregation-prone core (residues 42–48) within specific intermediates, implicating localized core exposure rather than wholesale unfolding as the trigger for misfolding. Temperature-dependent secondary-structure profiling confirms SAA_1−76_ behaves as a folded bundle with disordered loops. These findings highlight helix III stabilization and amyloidogenic segment masking as potential therapeutic strategies.

**TOC Graphic:** 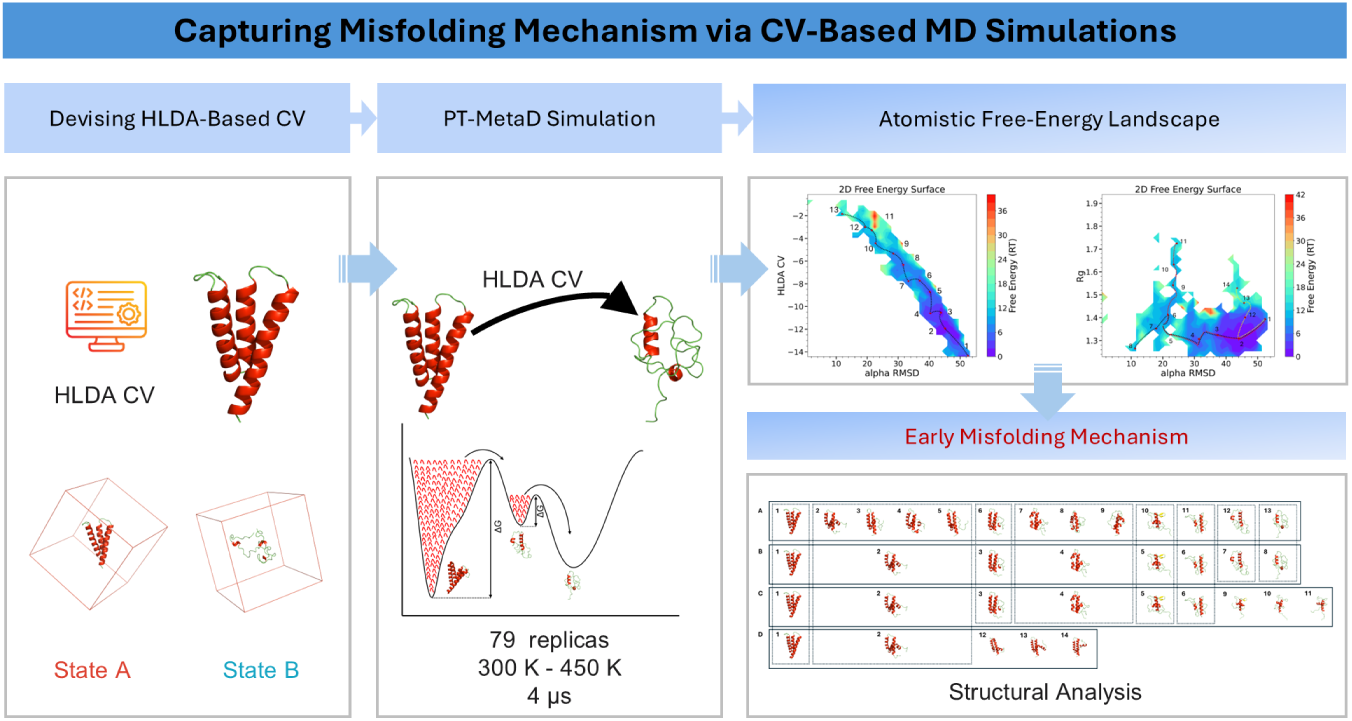

## Main

AA amyloidosis is a severe systemic disorder and a life-threatening complication of chronic inflammation, characterized by the deposition of insoluble amyloid fibrils in various organs and tissues, leading to progressive dysfunction. This condition arises as a long-term complication of chronic inflammatory diseases, including persistent infections, rheumatoid arthritis, and certain cancers.^1,2^ Recent findings also pointed out a link between alkaptonuria, an ultra-rare genetic disorder of tyrosine metabolism, persistent low-grade inflammation, and AA amyloidosis, ^3–5^ with a pro-aggregating effect of homogentisic acid (HGA) towards SAA highlighted in vitro.^6,7^ The principal component of AA amyloid deposits is the serum amyloid A (SAA) protein,^8^ a family of acute-phase apolipoproteins primarily produced by the liver in response to inflammatory stimuli. SAA proteins are normally associated with high-density lipoprotein (HDL) in circulation and play a critical role in modulating the immune response.^9^ SAA isoforms are acute-phase response proteins that are synthesized predominantly in the liver and expressed constitutively (constitutive SAA) or in response to inflammatory stimuli (acute phase SAA). SAA proteins are encoded by the SAA1, SAA2, SAA3, and SAA4 genes, located on chromosome 11p15.1 and expressed isoforms consist of 103-104 residues that are highly conserved throughout evolution. ^10^ SAA1 and SAA2 are the predominant acute-phase isoforms and their serum levels rise dramatically, up to 1000-fold, in response to infections, traumas, or other stimuli. ^11^ The exact biological function of SAA is only partly understood with increasing evidence suggesting SAA roles in cholesterol transport, antibody regulation, inhibition of platelet aggregation, and modulation of macrophage activity.^12^ Structural studies, including the crystallographic analysis of the native human Serum Amyloid A-1 (SAA1) (PDB ID 4IP9) at 2.5 resolution. ^9^ suggest that SAA1 (104 a.a.) has a four-helix bundle structure with a cone-shaped array, in which the N termini of helices 1 and 3 and the C termini of helices 2 and 4 are packed together. The residues 1-27, 32–47, 50–69, and 73–88 form helices 1, 2, 3 and 4, respectively.^9^ It has been reported that SAA helix bundle features a hydrophilic interior partially filled with water^13^ and *α*-helices 1 and 3 feature a strong amphipathic property with a hydrophobic face.^14^ The C-terminal tail forms multiple salt bridges and hydrogen bonds with the *α*-helices 1, 2 and 4 wrapping around the bundle plays a key role to stabilize the helix bundle structure.^9^ Amphipathic helices 1 and 3 form an elongated concave hydrophobic surface with a curvature radius complementary to that of HDL suggesting the possible HDL binding site.^1^ The formation of amyloid fibrils in AA amyloidosis occurs in the kidneys, spleen and liver,^15^ stemming from the misfolding of circulating SAA1 after significant increment of serum SAA1 up to 1000 fold reaching 1 mg mL^−1^ due to chronic inflammation.^11^

This process is thought to be a stepwise proteolytic processing from the full-length precursor SAA(1–104) to shorter fibrillar species. It begins with dissociation from HDL, followed by a specific proteolytic cleavage in the interdomain linker between residues 76 and 77, mediated by proteases such as cathepsin B, which generates the SAA(1–76) fragment.^16^ This 76-residue fragment is well-established as a highly unstable and profoundly amyloidogenic intermediate,^11,17,18^. It is this inherent instability, induced by the removal of the C-terminal stabilizing tail (residues 77-104). Subsequent proteolytic trimming to residues 67–69, as revealed by ex vivo structural studies^19,20^, likely occurs during or after this initial aggregation step, refining and stabilizing the end-stage fibril core. We therefore focus on SAA(1–76) in order to capture the crucial first step in the pathological cascade and to model the initial misfolding event.

In addition, by quantifying the hydrophobic residues and the aggregation-prone regions (APRs) exposure in intermediate conformational states, we investigate whether the aggregation propensity of SAA is driven primarily by its high concentration or by substantial exposure of APRs. Moreover, given the conflicting findings about the classification of SAA as an intrinsically disordered protein (IDP)^1^ or a folded protein with only partially disordered segments^9^, our work also aims to clarify this debate by providing an atomistic description of the early events in the misfolding of SAA along and place them on the free energy landscape of the monomeric protein.

To address these issues, we employ a supervised learning classification method (HLDA) to construct a collective variable for biasing simulations in PT-MetaD. This approach enables comprehensive exploration of the phase space of the system, providing free energy landscapes projected onto various variables and allowing us to characterize the structural ensembles corresponding to metastable states. Our main objective is to elucidate the early events in the misfolding of SAA, by probing the structural features of the identified metastable states and proposing a possible misfolding pathway.

We began our study by performing two 20 *µ*s unbiased trajectories for the folded and unfolded states of SAA (as can be seen in **Computational Methods**). Next, we calculated six selected descriptors that could potentially describe the misfolding process of SAA along the unbiased trajectories for both the folded and unfolded states (see **Computational Methods**). By applying HLDA and after the optimization, we selected the eigenvectors corresponding to the highest eigenvalue, as shown in **Table** 1. This choice ensures the maximum separation between the folded and unfolded states. The analysis of the weight distributions, illustrated in **Figure S1b**, provides structural insights into the system, indicating that the features of the folding process were captured by the CVs. Notably, the majority of the weight is attributed to the descriptors contact map-1 (CM-1), which represents the distance between *α*-Helix I and *α*-Helix II, followed by the *α*RMSD values of *α*-Helix III and *α*-Helix II. The optimized HLDA CV, namely the one with higher eigenvalue was defined as:

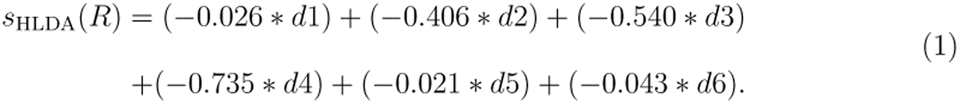

**Table 1:**
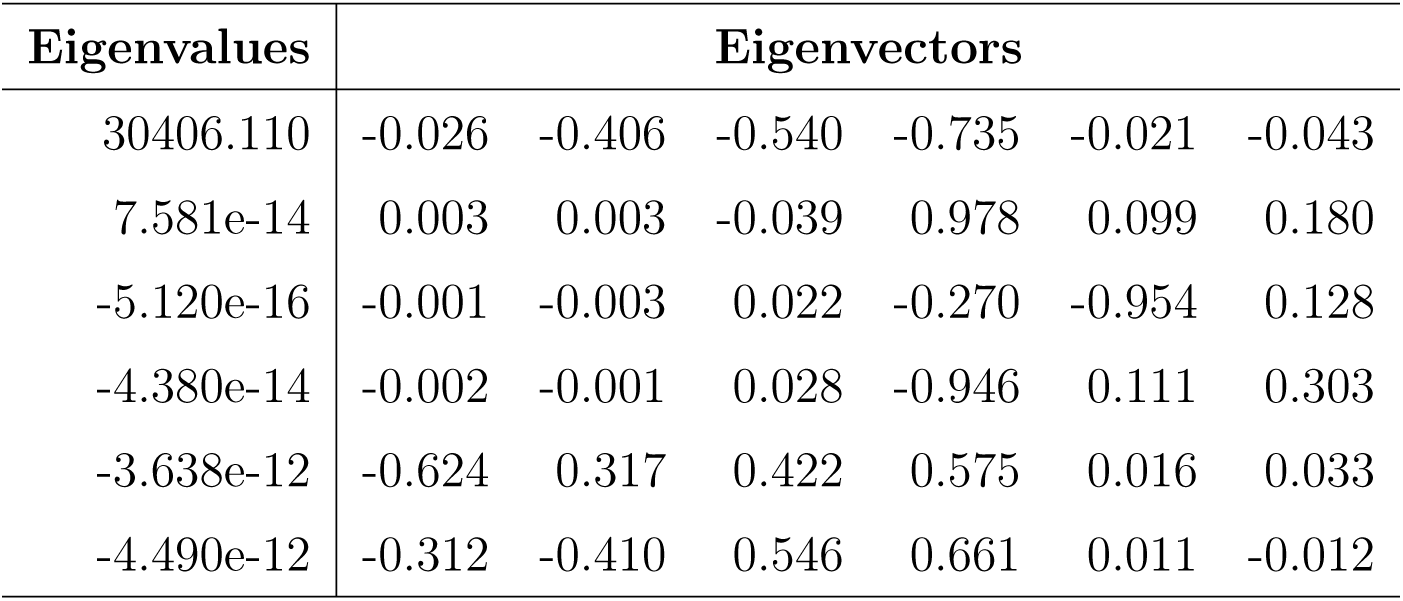
Eigenvalues and their corresponding eigenvectors.

Using the HLDA derived CV, **s**_HLDA_(**R**), we performed PT-MetaD simulations to explore the misfolding mechanism of SAA, in terms of different free energy landscape projections. The overlap of potential energy distributions across consecutive replicas, as illustrated in **Figure S2**, highlights the efficiency of the sampling and exchange rate between replicas. Additionally, the diffusion of the HLDA CV at T = 310 K over time (**Figure S3**) under the effect of the PT-MetaD potential demonstrates increased fluctuations reflecting enhanced exploration of phase space and improved sampling. The simulations convergence is confirmed through the effective diffusion of replica 06 (at a temperature of 310 K) across the temperature space (shown in **Figure S4**) along with the superposition of the time-dependent FES along various CVs (**Figures S5, S6, S7**) where the FES becomes static as a function of time.

From the MD simulations we obtained two 2D free energy landscapes projected on HLDA CV and *α*RMSD for the first one (**Figure 1a**), while for the second one is on *α*RMSD and radius of gyration (Rg) (**Figure 1b**).

**Figure 1:**
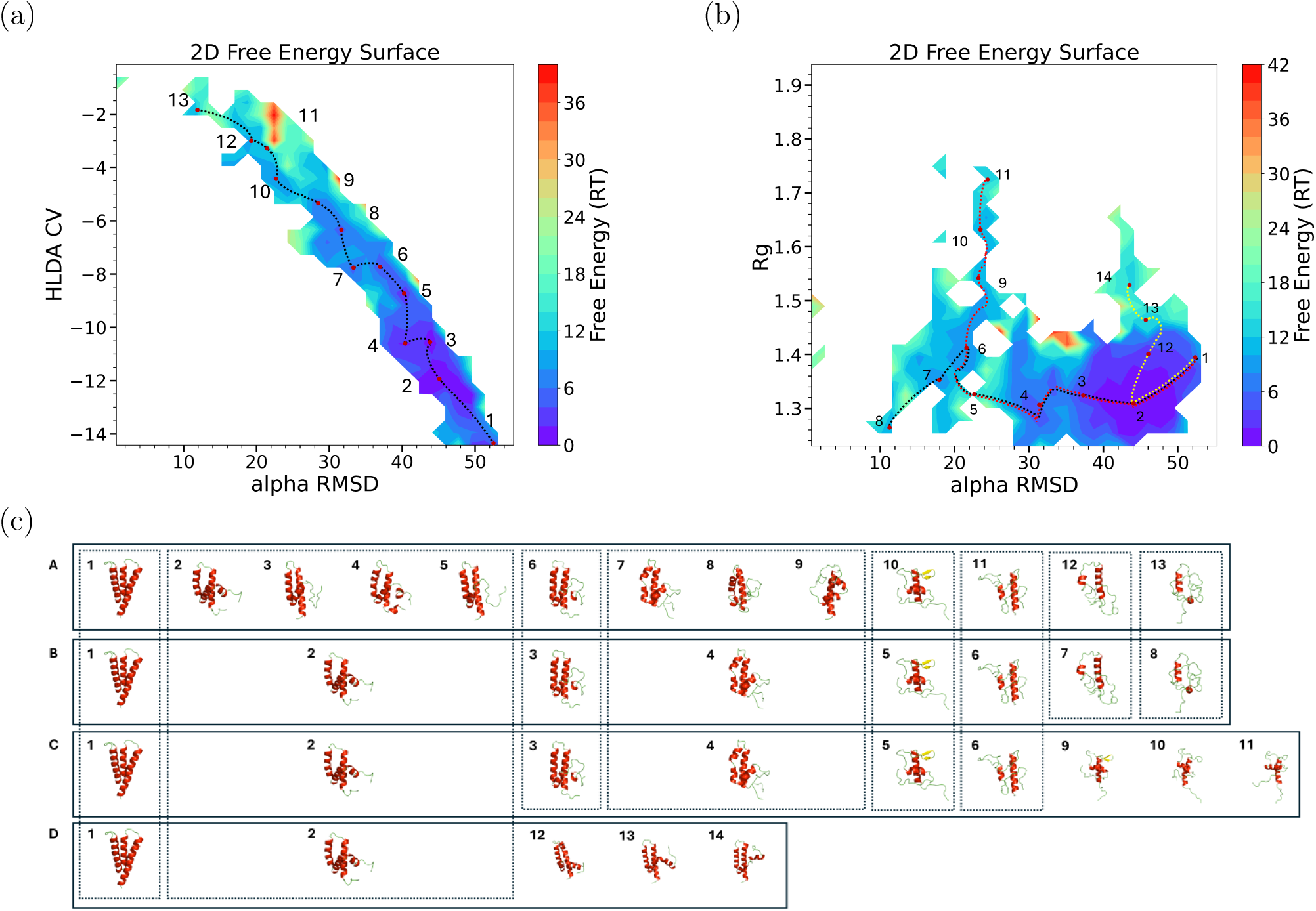
(a) FES as a function of *α*RMSD and HLDA showing a possible misfolding pathway. (b) FES as a function of the *α*RMSD and Rg showing three possible misfolding pathways and the 14 minima crossed by them. (c) A: Structures of the 13 minima crossed along the possible misfolding pathway shown in (a). B-D: Representative structures sampled during the simulations along the first, second and third possible misfolding pathways. Common structures found in shared basins among these pathways are enclosed within dotted boxes

The possible misfolding process is represented as a pathway connecting two basins: the initial folded structure and the final unfolded state, crossing a series of sampled different short-lived intermediates.

Using the MEPSA (Minimum Energy Path Surface Analysis) software^21^, we identified the minimum energy pathways as a function of collective variables we deemed informative on the misfolding mechanism. We would like to stress these pathways are neither kinetic pathways nor necessarily describe the committor function but are rather informative pathways along FES projections. These pathways connect between low free energy states, which correspond to the most probable configurations. The protein predominantly resides in these states, though it can still transition between them stochastically over time.

Thirteen distinct metastable states along the possible misfolding pathway were identified in the first FES (**Figure 1a**), labeled 1 to 13. Representative conformations from these basins are shown in panel A (**Figure 1c**), where the conformer in minimum 1 corresponds to a fully folded structure, while metastable state 13 represents an almost fully unfolded structure forming up to 25% of helical content with respect to the folded state structure.

To gain deeper insight into the structural changes along the possible misfolding pathway, we analyzed the *α*-helical content of the three *α*-helices in each of the thirteen minima illustrated in **Figure 2a**. The results indicate that *α*-Helix III unfolds first, followed by a sudden dissolution of *α*-Helix II after metastable state 5. The last helix to unfold is *α*-Helix I, which starts to decay after metastable state 6, with half of it disappearing by metastable state 13. The order of *α*-helix unfolding aligns with the weights of the descriptors shown in **Figure S1b**. As shown in **Figure 2b**, the overall *α*-helical content gradually decreases along the misfolding pathway, accompanied by an increase in random coil structure, ultimately reaching an almost fully unfolded state in metastable state 13. Rg analysis of structures in **Figure 2c** suggests that, despite the loss of secondary structure, the compactness of the fragment is not significantly affected. This observation is further supported by the contact map analysis (**Figure 2d**). Although there are fluctuations in the distance between the *α* helices, the distances between the centers of mass of the three *α*-helices remain relatively stable as we can notice in the metastable 13 with respect to the folded state structure.

**Figure 2:**
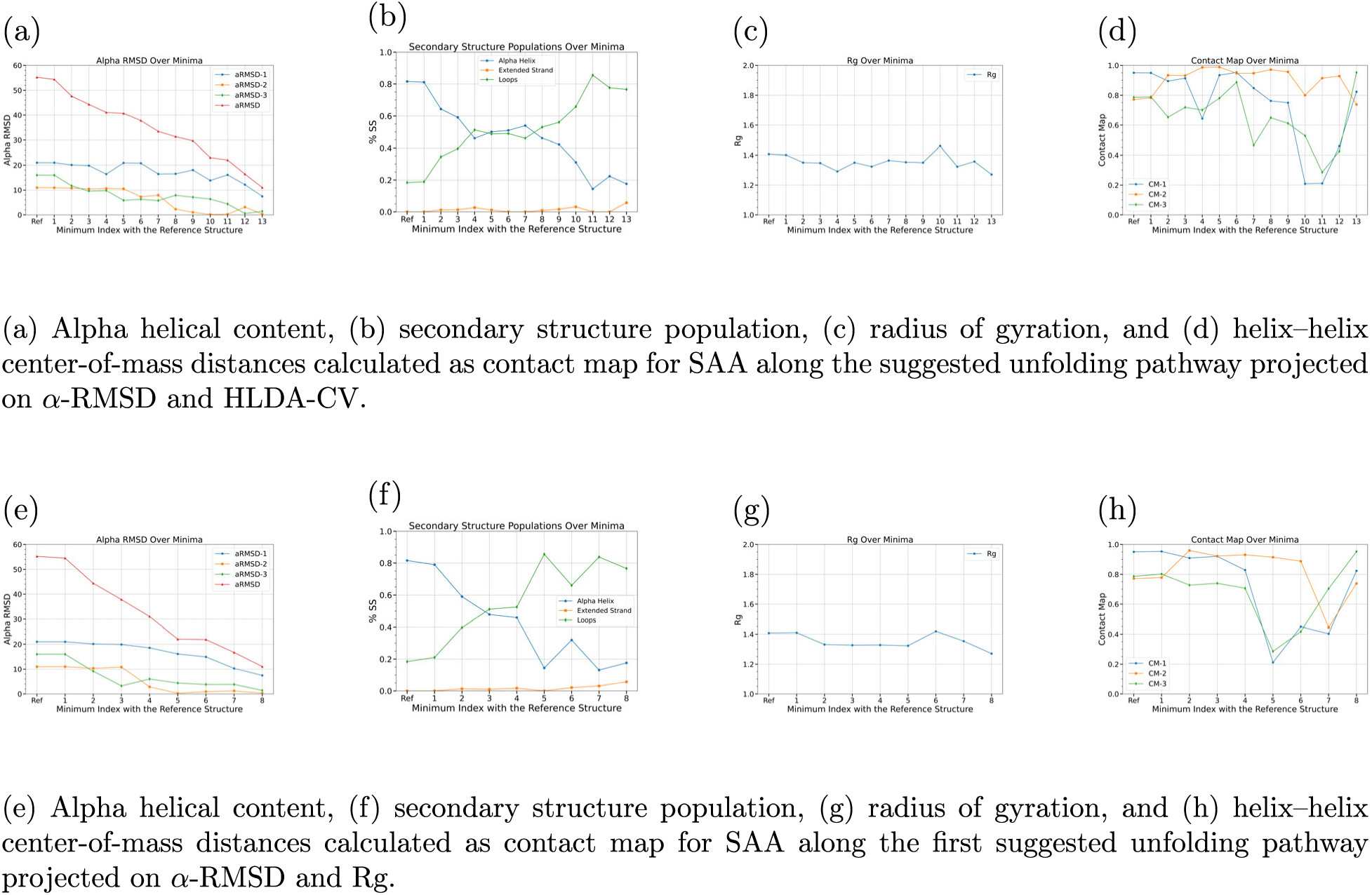
Structural analyses of SAA along (a-d) the suggested unfolding pathway projected on *α*-RMSD and HLDA-CV and (e-h) the first suggested unfolding pathway projected on *α*-RMSD and Rg, showing alpha helical content, secondary structure population, radius of gyration, and helix–helix center-of-mass distances for representative structures.

Along the misfolding pathway, we observe that the *α*-helices gradually transform into random coil in a specific sequence while maintaining structural compactness and preserving the tertiary structure. This process results in a structure composed almost entirely of random coil with some remaining *α*-helical content, yet the compactness remains intact.

These findings suggest a stepwise misfolding process, where *α*-helix III unfolds first followed by *α*-helix II and finally *α*-helix I while the overall compactness of SAA remains largely stable.

The solvent-accessible surface area (SASA) analysis (**Figure S8**) along the possible mis-folding pathway shows that the total SASA does not increase significantly along the unfolding pathway at the misfolded intermediate states. Similarly, the SASA of the hydrophobic residues (**Figure S8**) remain relatively stable during the misfolding. However, the SASA of the predicted aggregation-prone region (APR, corresponding to residues **42-48** as predicted by FuzDrop^22–24)^ (**Figure 3a**) remains relatively stable during the unfolding except the basins 9, 10, 11 and 12 in which the APR region becomes more exposed to the solvent.

**Figure 3:**
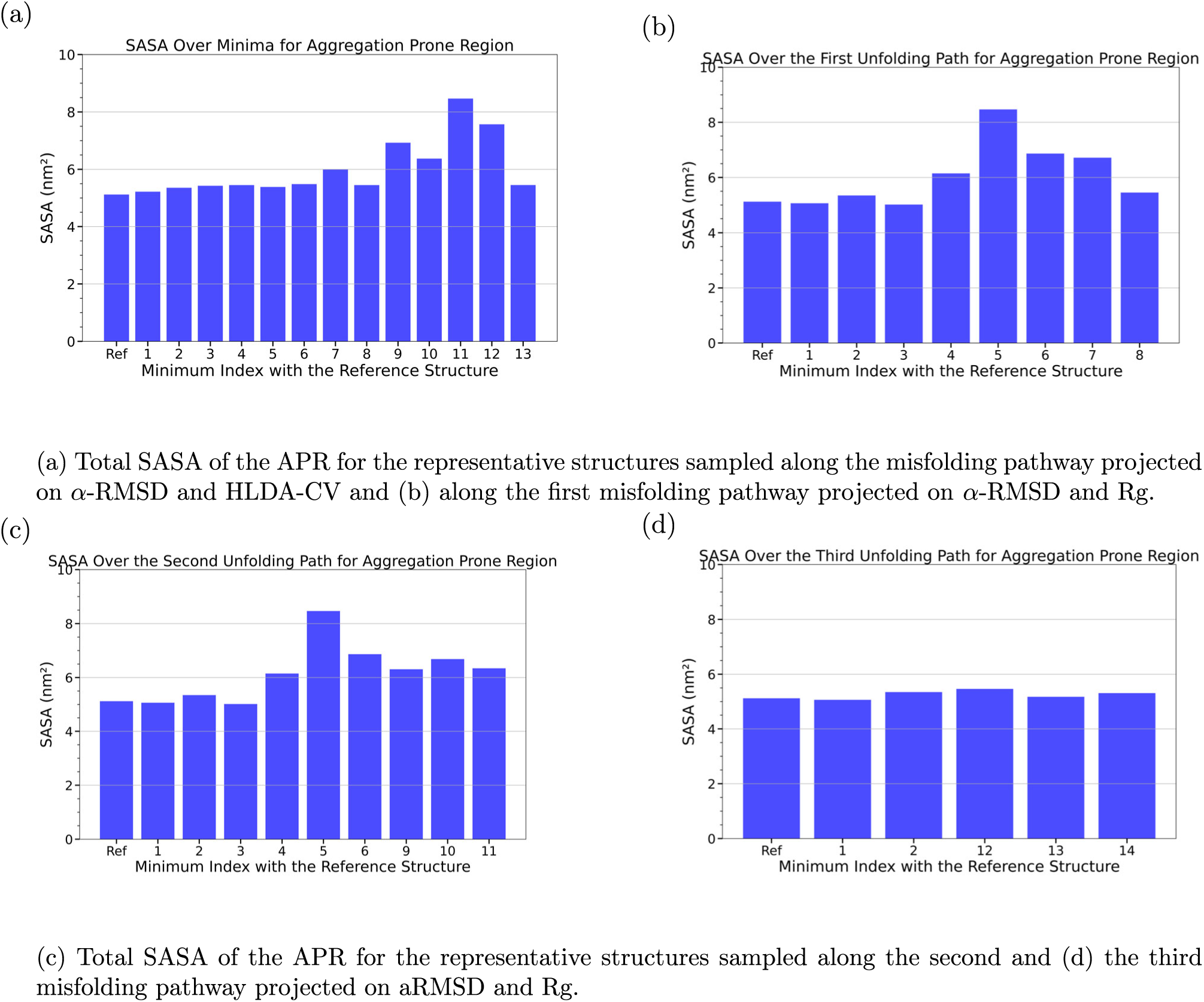
Total SASA of the APR for representative structures of SAA sampled along the distinct misfolding pathways: (a) projected on *α*-RMSD and HLDA-CV, (b-d) projected on *α*-RMSD and Rg.

These results suggest that the aggregation of SAA can be triggered by the increased solvent exposure of the APR of the aforementioned metastable states.

Figure 1b presents a 2D FES plotted as a function of *α*-RMSD and Rg, showing three possible misfolding pathways represented by black, red and yellow dotted lines, respectively. The first proposed pathway crosses eight basins, with the conformer in minimum 1 corresponding to a fully folded structure and basin 8 representing an almost fully unfolded structure as shown in Figure 1c, panel B. This pathway is aligned with the projected one along *α*-RMSD and HLDA-CV that has been discussed before, however, it is less tuned as we can see in Figure 1b. The basins 2, 3, 4 and 5 in the 2D FES along *α*-RMSD and HLDA-CV are gathered in basin 2 in the 2D FES along *α*-RMSD and Rg. Similarly, basins 7, 8 and 9 in the 2D FES along *α*-RMSD and HLDA-CV are gathered in basin 4 in the 2D FES along *α*-RMSD and Rg.

The results of the *α*-helicity analysis align with previous results, showing that *α*-Helix III unfolds first, followed by *α*-Helix II, and finally *α*-Helix I, as illustrated in Figure 2e. Along this pathway, the overall *α*-helical content decreases steadily while the random coil content increases, ending with a nearly fully unfolded structure in basin 8 as shown in Figure 2f. Despite the loss of secondary structure, the compactness of the protein remains stable throughout the pathway, as revealed by the Rg analysis in Figure 2g. The contact map analysis further supports this, demonstrating that the distances between the centers of mass of the *α*-helices remain relatively unchanged in basin 8 compared to basin 1. However, in metastable states 5, 6 and 7, there is a marked increase in the distances between the *α* helices which align with the contact map analysis of the 2D FES along *α*RMSD and HLDA-CV where we can notice the increase of distances in basins 10, 11, and 12 (Figure 2d) which are the same metastable states in both pathways (Figure 1c).

These results align with the structural analyses of the previously discussed misfolding pathway projected on *α*RMSD and HLDA CV. It corroborates the stochastic switch between the folded and misfolded configurations passing through different intermediates in a sequential process starting by the destabilization of *α* helix III.

SASA analysis reveals that the overall solvent-exposure area and the SASA attributed to hydrophobic residues remains largely unchanged across the misfolded metastable states (**Figure S9**). However, the SASA attributed to the APR increased significantly during unfolding in the basins 4, 5, 6 and 7 (Figure 3b) which are the same basins 9, 10, 11 and 12 in the 2D FES along *α*RMSD and HLDA-CV that show a significant increase of SASA attributed to APR (Figure 3a). These findings support the hypothesis that the localized core exposure, namely the APR (residues 42-48), within specific misfolded conformers may trigger the aggregation process.

Regarding the energy barriers along the pathway identified by MEPSA, the analysis reveals a series of metastable states separated by barriers ranging from ∼2 RT to ∼6 RT as we can see in Figure 1. The highest barrier (∼6 RT) corresponds to the initial step out of the native basin, which is likely the rate-limiting step for the initiation of misfolding under physiological conditions. Subsequent transitions between intermediates involve smaller barriers (∼2-4 RT), suggesting that once the native fold is destabilized, the protein can sample the misfolded intermediates with relatively higher probability. It is important to note that these barriers are derived from projections of the high-dimensional free energy landscape onto one or two collective variables. While they provide a valuable thermodynamic perspective on the relative stability of states, their absolute heights should be interpreted with caution regarding kinetics. The committor probabilities and exact transition rates would require dedicated reaction coordinate analysis and extensive transition path sampling, which is beyond the scope of this current study but represents a compelling direction for future work.

The second identified pathway represented by red dotted line in Figure 1b crosses nine metastable states but does not lead to a fully unfolded structure. The conformer in metastable state 11 remains partially unfolded, as shown in Figure 1c Panel C sharing the first 6 basins with the first pathway.

The *α*-helicity analysis reveals that *α*-Helix III unfolds completely first, followed by *α*-Helix II, and finally *α*-Helix I within the first six metastable states, which overlap with the initial part of the first suggested pathway. However, in the rest three basins (9, 10 and 11), *α*-Helix I remains stable while *α*-Helix-II increases slightly as we can see in Figure 4a. Additionally, Figure 4b shows a steady decrease in overall *α*-helical content and a corresponding increase in random coil content up to basin 5, after which a significant increase occurs in the last four basins ending with a structure forming up to 50% of helical content with respect to the folded state structure.

**Figure 4:**
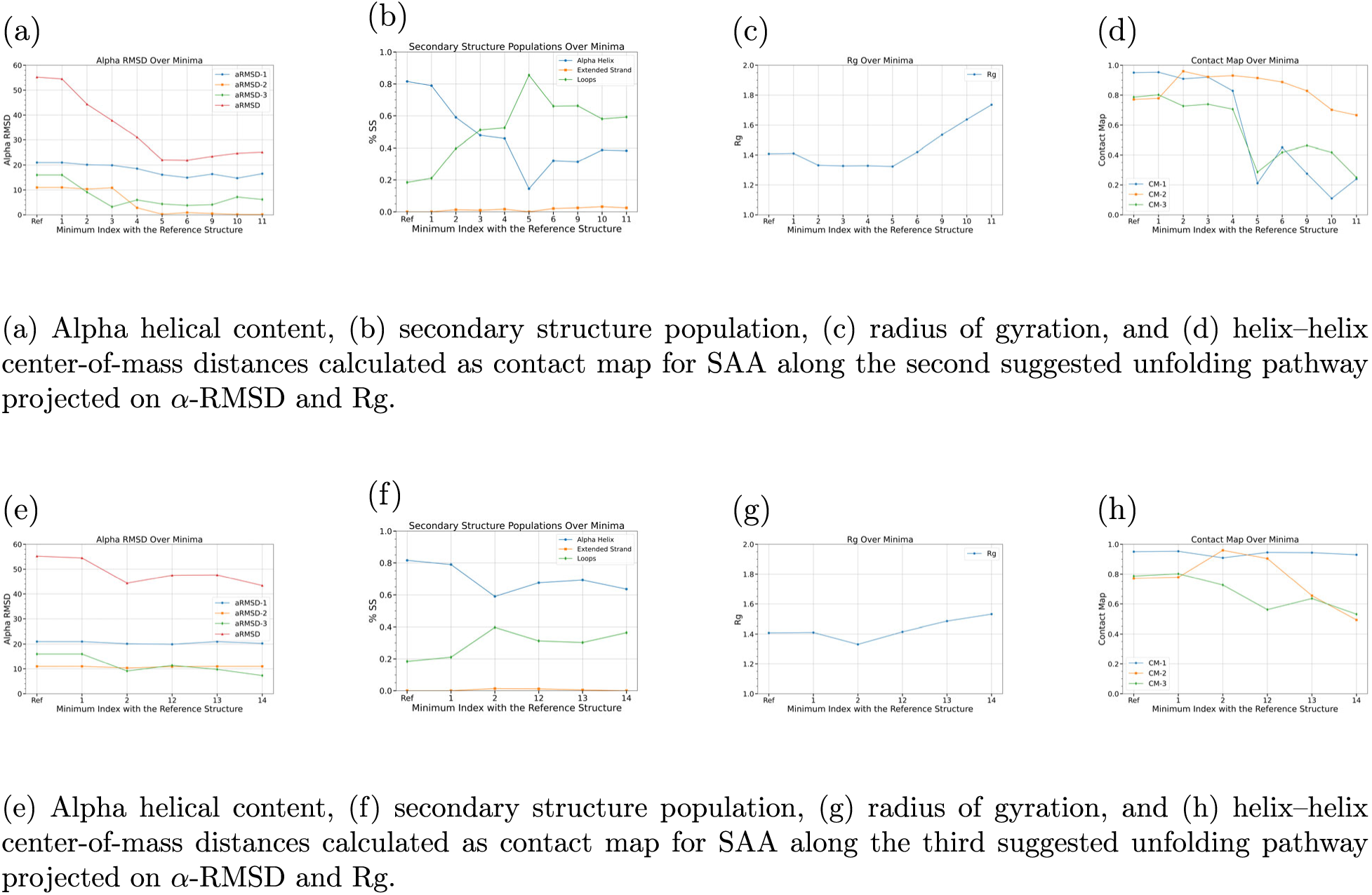
Structural analyses of SAA along (a-d) the second and (e-h) the third suggested unfolding pathway projected on *α*-RMSD and Rg, showing alpha helical content, secondary structure population, radius of gyration, and helix–helix center-of-mass distances for representative structures.

The Rg analysis indicates a significant increase in the last four metastable states, signifying a loss of protein compactness, as shown in Figure 4c. These findings are supported by the contact map analysis (Figure 4d), which reveals a significant increase in the distances between the *α*-helices, particularly between *α*-Helix I and II and between *α*-Helix II and III. Similar observations are noticed in the third suggested pathway represented by yellow dotted line in Figure 1b, which passes from metastable state 1 to metastable state 14. In this pathway, the *α*-helices I and II were stable while the *α*-helix III content decreases slightly (Figure 4e) ending with a structure that has around 80% of helical content with respect to the folded state structure as shown in Figure 4f. However, the protein’s compactness decreases, as evidenced by an increase in the Rg value (Figure 4g), and the distance between *α*-Helix I and III and between *α*-Helix II and III also increase as illustrated in Figure 4h.

The final two off-pathways represent a different misfolding mechanism, characterized by an initial loss of the compactness while the secondary structure elements remain relatively intact. However, for the analyzed SAA fragment, this mechanism appears improbable because the pathways fail to reach a fully unfolded structure. This observation further corroborates the conclusion that destabilization of *α*-helix III drives the misfolding process.

Additionally, the total SASA (**Figure S10**) and the SASA of the APR (Figure 3c) increase along the second pathway suggesting that the increased exposure of the APR drives the aggregation process. In contrast, pathway 3 shows no significant change in total SASA and hydrophobic SASA (**Figure S11**), or APR SASA (Figure 3d). However, since these two off-pathways do not lead to complete unfolding, these results assert once again the hypothesis that SAA aggregation process is driven by the APR solvent exposure.

Wang et al.^25^ examined SAA(1–104) and fragments by molecular dynamics simulations, proposing two SAA(1–76) motifs (‘helix-weakened’ and ‘helix-broken’) in which helix III re-aligns relative to helix I and modulates the accessibility of the N-terminal segment. In contrast, our HLDA-guided PT-MetaD simulations yield converged free-energy surfaces and pathways that resolve thirteen intermediates and a sequential order of structural changes: helix III destabilizes first, then helix II and helix I, while overall compactness is retained. This state-resolved landscape further identifies transient exposure of residues 42-48 as the local trigger of misfolding, providing a mechanistic basis for targeted stabilization/masking strategies.

There have been controversial experimental findings about the nature of SAA as IDP^1^ or folded protein with only partially disordered segments. ^9^ To resolve this debate, temperature-dependent secondary structure analysis of SAA_1−76_ structural ensembles, as shown in Figure 5 has been performed. The high temperatures (up to 450 K) employed in our PT-MetaD protocol are indeed unphysical; however, they are used solely as a computational tool to accelerate sampling and overcome the high free-energy barriers associated with protein folding on simulation timescales. The replicas at these high temperatures are essential for ensuring efficient random walks in temperature space and facilitating exchanges that prevent the low-temperature (physiological) replicas from becoming trapped in local minima. The purpose of this analysis was not to characterize physiological behavior at high temperature, but to conduct a controlled computational experiment to test a hypothesis about the classification of SAA. IDPs often exhibit a gain of structure (folding) at higher temperatures due to the strengthening of hydrophobic interactions. Our observation that SAA(1-76) becomes increasingly disordered (random coil) with rising temperature is a classic signature of a folded protein that denatures upon heating, not an IDP. This contrast helps resolve the debate referenced in our introduction and strengthens our argument that SAA is a folded bundle with flexible loops.

**Figure 5:**
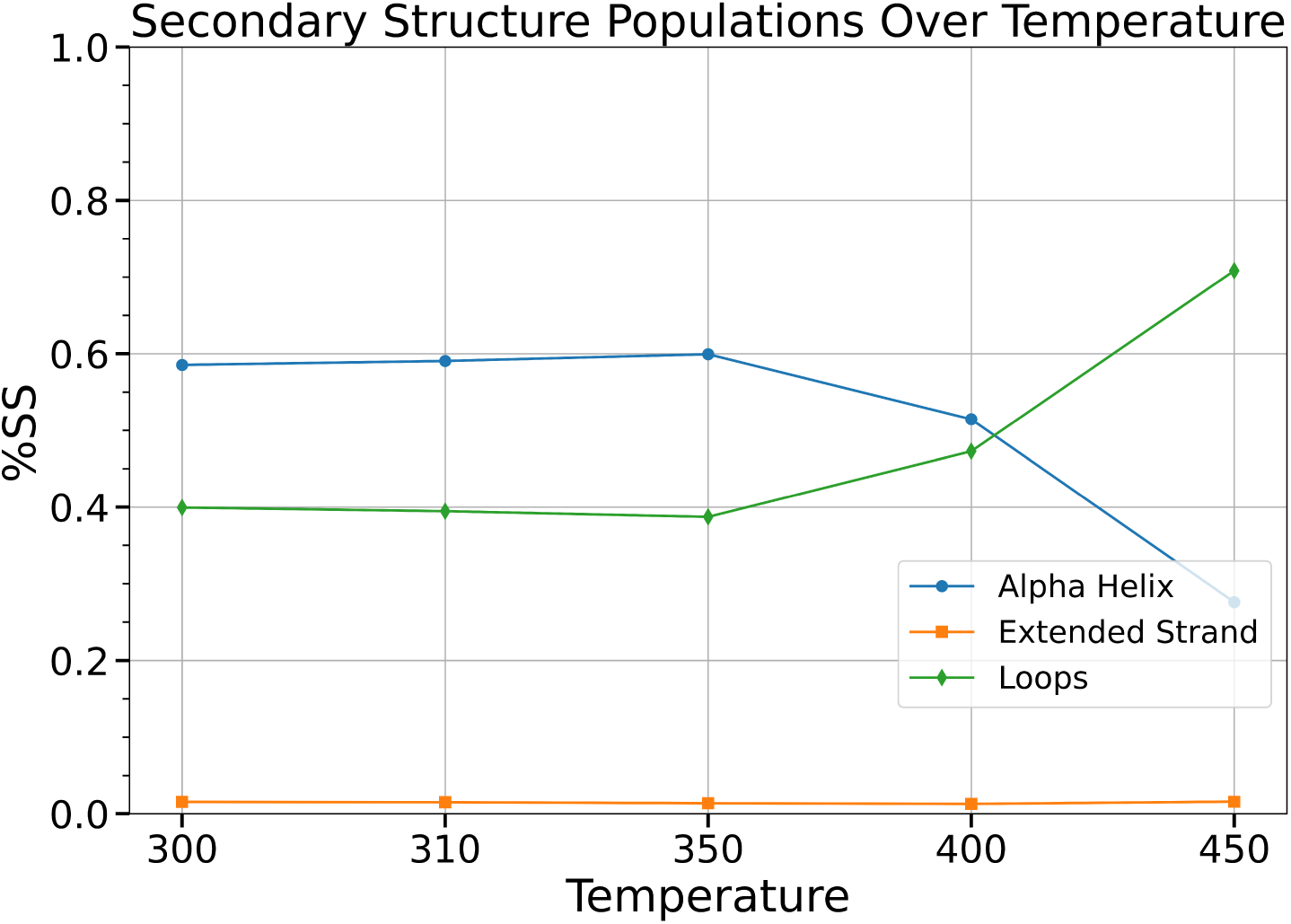
The secondary structure analysis of SAA over different temperatures. The high-temperature simulations in this analysis are a sampling device, and the secondary structure analysis at these temperatures is interpreted specifically in the context of assessing intrinsic stability and distinguishing the behavior of SAA from that of IDPs.

In this study, we explored the early misfolding steps of the pro-amyloidogenic 1-76 fragment of SAA (SAA_1−76_), which is the principal component of pathogenic fibrils, by combining a range of molecular dynamics simulations methods.

By implementing Harmonic Linear Discriminant Analysis (HLDA) to construct collective variables (CVs) and utilizing Parallel-Tempering Metadynamics (PT-MetaD), we mapped the free energy landscapes of the monomeric state of SAA_1−76_, identifying key metastable states along its possible misfolding pathway. Notably, by utilizing HLDA with a priori set of descriptors, we generated a linear collective variable that drives the PT-MetaD simulation toward the unfolded state providing free energy estimates. Furthermore, the weight distributions of the calculated HLDA CV reveals useful information, offering physical insight into the misfolding process. The simulations revealed a sequential misfolding process, where *α*-helix III unfolds first, followed by *α*-helix II and finally *α*-helix I. Despite the loss of secondary structure, the overall compactness of SAA remained largely intact. Moreover, SASA analyses show that the overall SASA, as well as that of hydrophobic residues remain relatively constant along the misfolding pathway at the misfolded intermediate states. However, the SASA of the predicted APR (residues 42–48) increases in a specific metastable states. This observation implies that SAA aggregation can be driven by the significant increase in the solvent exposure of APRs. Furthermore, the secondary structure analysis of SAA across different temperatures reveals that, contrary to the typical behavior of IDPs, which tend to become more ordered at elevated temperatures, SAA shows predominantly unstructured, random coil conformations at higher temperatures, while more structured conformations are evident at lower temperatures.

These results provide insights into the molecular basis of SAA misfolding and identify metastable states that could serve as potential targets for therapeutic intervention, such as strategies aim at stabilizing native *α*-helical regions. The approach presented here enhances our understanding of SAA aggregation and offers a framework for studying the early misfolding events of other amyloidogenic proteins. Future research should focus on experimental validation of these findings and on exploring small-molecule misfolding inhibitors, known as pharmacological chaperones^26,27^, that stabilize the native structure of SAA to mitigate its pathogenic aggregation.

## Computational Methods

### Molecular Dynamics (MD)

The crystallographic structure of SAA with code 4IP9 in the Protein Data Bank (PDB) was used to derive the fragment SAA_1−76_ as a starting configuration after deleting residues 77-104. ^9^ This truncated SAA form is the most common in pathological amyloid deposits, due to the pro-amyloidogenic destabilization of the four helix bundle after removal of the C-terminal residues. .^8,9,16^ All MD simulations were performed using the charmm36 force field^28^ and GROMACS 2022.3 software package. ^29^ The protein was solvated in TIP3P water^30^ and counter ions and additional salt ions (sodium and chloride) were added to neutralize the system and get a final salt concentration of 0.15 M. The system was then energy-minimized using the steepest descent method, with the simulations successfully terminating upon fulfill the energy criteria of a maximum force less than 1000 kJ/mol/nm. Followingly, an NPT equilibration at 310 K and 1 atm was performed for 100 ps, using the V-rescale thermostat^31^ and Parrinello-Rahman barostat ^32^. The particle-mesh Ewald method^33^ was used for long-range electrostatics with a short-range cutoff of 1.0 nm. A cutoff of 1.0 nm was used for the Lennard-Jones interactions. All bonds were constrained to their equilibrium length with the LINCS algorithm^34^. A 2 fs timestep was used. Finally, two 20 ns NVT simulations were performed at 310 K and 500 K respectively.

### Devising Collective Variables

According to Ref. 35, the Harmonic Linear Discriminant Analysis (HLDA) has been used to construct a 1-D collective variable (CV) that can describe the SAA folding process. This is a modification of Fisher’s linear discriminant analysis (LDA) that estimates the optimal dimensional projection *W* to achieve maximum separation for the unbiased distributions of the folded and unfolded states within an N*_d_*-dimensional descriptor space. In HLDA, the optimization of *W* is performed by maximizing the ratio of the so-called between-class scatter (*S_b_*) to the within-class scatter (*S_w_*), expressed as:

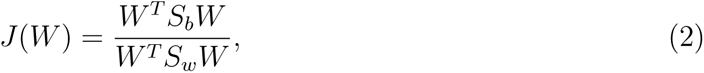

where *S_b_*quantifies the separation of the mean values of the two classes (folded and unfolded states) and is defined as:

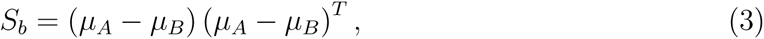

with *µ_A_*and *µ_B_* representing the mean values of the descriptors for the folded and unfolded states, respectively.

In contrast, *S_w_* measures the spread within each class. HLDA employs the harmonic average of the covariance matrices, defined as:

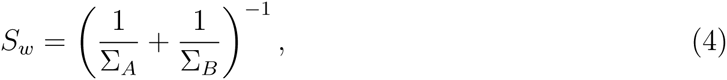

where Σ*_A_* and Σ*_B_* are the covariance matrices of the folded and unfolded states. Substituting these definitions into **Equation** 2, the optimization problem becomes the maximization of *J*(*W*):

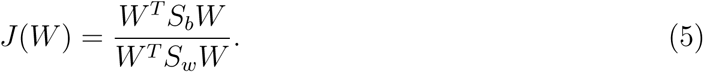

The optimal direction *W* ^∗^ is then determined by solving the eigenvalue problem:

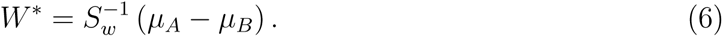

The resulting collective variable, denoted as **s**_HLDA_(**R**), is computed as the projection of the descriptors *d*(*R*) along the optimal direction *W* ^∗^, corresponding to the highest eigenvalue in **Equation** 6, formulated as:

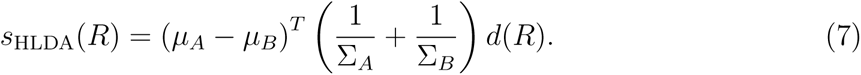

This approach is able to systematically construct a CV capable of distinguishing between folded and unfolded states, providing a one-dimensional representation of the complex folding dynamics.

Based on our intuition, we chose a set of six descriptors which can potentially describe the misfolding process of SAA. The first set probes the alpha helical content of each *α*-helix of SAA (d1, d2, d3) (*α*RMSD-1, *α*RMSD-2, and *α*RMSD-3). The second set consisted of three distances between the centers of each *α*-helix calculated and transformed by a switching function to a contact map (d4, d5, d6) (CM-1, CM-2, and CM-3) as shown in **Figure S1a**. The descriptors were calculated along the folded and unfolded state unbiased trajectories using the open-source, community-developed PLUMED library.^36–39^

By applying HLDA, we estimated the hyperplanes that best distinguished the unbiased distributions of the folded and unfolded states within the space defined by each set of descriptors constructing a collective variables as linear combinations of the set of descriptors.

### Parallel-Tempering Metadynamics (PTMetaD) Simulations

PTMetaD simulations^40^ were performed in the NPT ensemble by starting from 79 equidistant time-interval frames from the 20 ns folded state NPT MD obtained in the previous step. We ran 79 replicas in a temperature range between 300 K to 450 K for 50 ns each and cumulative simulation time of 3.9 *µ*s. Each replica was biased on the previously defined CV. The Gaussian height and width were 3.0 KJ/mol and 0.02, respectively, and a bias factor of 8 and a Gaussian deposition rate of 100 were used in all simulations. The MD and force field parameters were the same as in the unbiased NPT MD.

## Supporting Information

The Supporting Information is available free of charge at https://pubs.acs.org.

The six selected descriptors and their analysis of the weight distributions are shown in **Figure** S1. The overlap of potential energy probability distributions across consecutive replicas is illustrated in **Figure** S2. The diffusion of the HLDA CV at T = 310 K over time is demonstrated in **Figure** S3. The effective diffusion of replica 06 (at a temperature of 310 K) across the temperature space is shown in **Figure** S4. The superposition of the time-dependent FES along *α*RMSD, HLDA CV and Rg are illustrated in **Figures** S5, S6, S7, respectively. Total SASA for the protein and for the hydrophobic residues for the representative structure sampled during the simulations along *α*RMSD and HLDA CV is shown in **Figures** S8. Total SASA for the protein and for the hydrophobic residues for the representative structure sampled during the simulations along the first, second and third misfolding path along *α*RMSD and Rg are illustrated in **Figures** S9, S10, S11, respectively.

## Supporting information

The Supporting Information is available free of charge at https://pubs.acs.org.

## Acknowledgment

H.N acknowledges the Italian Ministry of University and Research (MUR) for a “Bando PON 37 ciclo - REACT-EU FSE DM 1061” grant. H.N acknowledges the CINECA award under the ISCRA initiative for the availability of high-performance computing resources and support. D.B. acknowledges support from the University of Siena (grant PSR 2021 F-CUR).

## References

(1) Frame, N. M.; Gursky, O. Structure of serum amyloid A suggests a mechanism for selective lipoprotein binding and functions: SAA as a hub in macromolecular interaction networks. FEBS letters 2016, 590, 866–879.

(2) Pettersson, T.; Konttinen, Y. T.; Maury, C. P. J. Treatment strategies for amyloid A amyloidosis. Expert opinion on pharmacotherapy 2008, 9, 2117–2128.

(3) Bernini, A.; Spiga, O.; Santucci, A. Structure-function relationship of homogentisate 1, 2-dioxygenase: understanding the genotype-phenotype correlations in the rare genetic disease alkaptonuria. Current Protein and Peptide Science 2023, *24*, 380–392.

(4) Braconi, D.; Geminiani, M.; Psarelli, E. E.; Giustarini, D.; Marzocchi, B.; Rossi, R.; Bernardini, G.; Spiga, O.; Gallagher, J. A.; Le Quan Sang, K.-H.; Arnoux, J.-B.; Imrich, R.; Al-Sbou, M. S.; Gornall, M.; Jackson, R.; Ranganath, L. R.; Santucci, A. Effects of nitisinone on oxidative and inflammatory markers in alkaptonuria: results from SONIA1 and SONIA2 studies. Cells 2022, 11, 3668.

(5) Braconi, D.; Giustarini, D.; Marzocchi, B.; Peruzzi, L.; Margollicci, M.; Rossi, R.; Bernardini, G.; Millucci, L.; Gallagher, J.; Le Quan Sang, K.-H.; Imrich, R.; Rovensky, J.; Al-Sbou, M.; Ranganath, L.; Santucci, A. Inflammatory and oxidative stress biomarkers in alkaptonuria: data from the DevelopAKUre project. Osteoarthritis and cartilage 2018, 26, 1078–1086.

(6) Braconi, D.; Millucci, L.; Bernini, A.; Spiga, O.; Lupetti, P.; Marzocchi, B.; Niccolai, N.; Bernardini, G.; Santucci, A. Homogentisic acid induces aggregation and fibrillation of amyloidogenic proteins. Biochimica et Biophysica Acta (BBA)-General Subjects 2017, 1861, 135–146.

(7) Mastroeni, P.; Trezza, A.; Geminiani, M.; Frusciante, L.; Visibelli, A.; Santucci, A. HGA Triggers SAA Aggregation and Accelerates Fibril Formation in the C20/A4 Alkaptonuria Cell Model. Cells 2024, 13, 1501.

(8) Sack Jr, G. H. Serum amyloid A–a review. Molecular medicine 2018, 24, 46.

(9) Lu, J.; Yu, Y.; Zhu, I.; Cheng, Y.; Sun, P. D. Structural mechanism of serum amyloid A-mediated inflammatory amyloidosis. Proceedings of the National Academy of Sciences 2014, 111, 5189–5194.

(10) Sun, L.; Richard, D. Y. Serum amyloid A1: structure, function and gene polymorphism. Gene 2016, 583, 48–57.

(11) Westermark, G. T.; Fändrich, M.; Westermark, P. AA amyloidosis: pathogenesis and targeted therapy. Annual Review of Pathology: Mechanisms of Disease 2015, 10, 321–344.

(12) Kisilevsky, R.; Manley, P. N. Acute-phase serum amyloid A: perspectives on its physiological and pathological roles. Amyloid 2012, 19, 5–14.

(13) Smole, U.; Kratzer, B.; Pickl, W. F. Soluble pattern recognition molecules: guardians and regulators of homeostasis at airway mucosal surfaces. European journal of immunology 2020, 50, 624–642.

(14) Hu, Z.; Bang, Y.-J.; Ruhn, K. A.; Hooper, L. V. Molecular basis for retinol binding by serum amyloid A during infection. Proceedings of the National Academy of Sciences 2019, 116, 19077–19082.

(15) Rennegarbe, M.; Lenter, I.; Schierhorn, A.; Sawilla, R.; Haupt, C. Influence of C-terminal truncation of murine Serum amyloid A on fibril structure. Scientific Reports 2017, 7, 6170.

(16) Yamada, T.; Liepnieks, J.; Kluve-Beckerman, B.; Benson, M. Cathepsin B generates the most common form of amyloid A (76 residues) as a degradation product from serum amyloid A. Scandinavian journal of immunology 1995, 41, 94–97.

(17) Uhlar, C. M.; Whitehead, A. S. Serum amyloid A, the major vertebrate acute-phase reactant. European journal of biochemistry 1999, 265, 501–523.

(18) Simons, J. P.; Al-Shawi, R.; Ellmerich, S.; Speck, I.; Aslam, S.; Hutchinson, W. L.; Mangione, P. P.; Disterer, P.; Gilbertson, J. A.; Hunt, T.; Millar, D. J.; Minogue, S.; Bodin, K.; Pepys, M. B.; Hawkins, P. N. Pathogenetic mechanisms of amyloid A amyloidosis. Proceedings of the National Academy of Sciences 2013, 110, 16115–16120.

(19) Liberta, F.; Loerch, S.; Rennegarbe, M.; Schierhorn, A.; Westermark, P.; Westermark, G. T.; Hazenberg, B. P.; Grigorieff, N.; Fändrich, M.; Schmidt, M. Cryo-EM fibril structures from systemic AA amyloidosis reveal the species complementarity of pathological amyloids. Nature communications 2019, 10, 1104.

(20) Banerjee, S.; Baur, J.; Daniel, C.; Pfeiffer, P. B.; Hitzenberger, M.; Kuhn, L.; Wiese, S.; Bijzet, J.; Haupt, C.; Amann, K. U.; Zacharias, M.; Hazenberg, B. P. C.; Westermark, G. T.; Schmidt, M.; Fändrich, M. Amyloid fibril structure from the vascular variant of systemic AA amyloidosis. Nature Communications 2022, 13, 7261.

(21) Marcos-Alcalde, I.; Setoain, J.; Mendieta-Moreno, J. I.; Mendieta, J.; Gomez-Puertas, P. MEPSA: minimum energy pathway analysis for energy landscapes. Bioinformatics 2015, 31, 3853–3855.

(22) Hardenberg, M.; Horvath, A.; Ambrus, V.; Fuxreiter, M.; Vendruscolo, M. Widespread occurrence of the droplet state of proteins in the human proteome. Proceedings of the National Academy of Sciences 2020, 117, 33254–33262.

(23) Vendruscolo, M.; Fuxreiter, M. Sequence determinants of the aggregation of proteins within condensates generated by liquid-liquid phase separation. Journal of Molecular Biology 2022, 434, 167201.

(24) Hatos, A.; Tosatto, S. C.; Vendruscolo, M.; Fuxreiter, M. FuzDrop on AlphaFold: visualizing the sequence-dependent propensity of liquid–liquid phase separation and aggregation of proteins. Nucleic acids research 2022, 50, W337–W344.

(25) Wang, W.; Khatua, P.; Hansmann, U. H. Cleavage, downregulation, and aggregation of serum amyloid A. The Journal of Physical Chemistry B 2020, 124, 1009–1019.

(26) Cohen, F. E.; Kelly, J. W. Therapeutic approaches to protein-misfolding diseases. Nature 2003, 426, 905–909.

(27) Vendruscolo, M. The thermodynamic hypothesis of protein aggregation. Molecular Aspects of Medicine 2025, 103, 101364.

(28) Best, R. B.; Zhu, X.; Shim, J.; Lopes, P. E.; Mittal, J.; Feig, M.; MacKerell Jr, A. D. Optimization of the additive CHARMM all-atom protein force field targeting improved sampling of the backbone *ϕ*, *ψ* and side-chain *χ*1 and *χ*2 dihedral angles. Journal of chemical theory and computation 2012, *8*, 3257–3273.

(29) Páll, S.; Abraham, M. J.; Kutzner, C.; Hess, B.; Lindahl, E. Tackling exascale software challenges in molecular dynamics simulations with GROMACS. Solving Software Challenges for Exascale: International Conference on Exascale Applications and Software, EASC 2014, Stockholm, Sweden, April 2-3, 2014, Revised Selected Papers 2. 2015; pp 3–27.

(30) Jorgensen, W. L.; Chandrasekhar, J.; Madura, J. D.; Impey, R. W.; Klein, M. L. Comparison of simple potential functions for simulating liquid water. The Journal of chemical physics 1983, 79, 926–935.

(31) Bussi, G.; Donadio, D.; Parrinello, M. Canonical sampling through velocity rescaling. The Journal of chemical physics 2007, 126, 014101.

(32) Parrinello, M.; Rahman, A. Polymorphic transitions in single crystals: A new molecular dynamics method. Journal of Applied physics 1981, 52, 7182–7190.

(33) Darden, T.; York, D.; Pedersen, L. Particle mesh Ewald: An N log (N) method for Ewald sums in large systems. The Journal of chemical physics 1993, 98, 10089–10092.

(34) Hess, B. P-LINCS: A parallel linear constraint solver for molecular simulation. Journal of chemical theory and computation 2008, 4, 116–122.

(35) Mendels, D.; Piccini, G.; Brotzakis, Z. F.; Yang, Y. I.; Parrinello, M. Folding a small protein using harmonic linear discriminant analysis. The Journal of chemical physics 2018, 149, 194113.

(36) Bonomi, M.; Branduardi, D.; Bussi, G.; Camilloni, C.; Provasi, D.; Raiteri, P.; Donadio, D.; Marinelli, F.; Pietrucci, F.; Broglia, R. A.; Parrinello, M. PLUMED: A portable plugin for free-energy calculations with molecular dynamics. Computer Physics Communications 2009, 180, 1961–1972.

(37) Tribello, G. A.; Bonomi, M.; Branduardi, D.; Camilloni, C.; Bussi, G. PLUMED 2: New feathers for an old bird. Computer physics communications 2014, 185, 604–613.

(38) Promoting transparency and reproducibility in enhanced molecular simulations. *Nature methods* 2019, *16*, 670–673.

(39) Tribello, G. A.; Bonomi, M.; Bussi, G.; Camilloni, C.; Armstrong, B. I.; Arsiccio, A.; Aureli, S.; Ballabio, F.; Bernetti, M.; Bonati, L.; Brookes, S. G. H.; Brotzakis, Z. F.; Capelli, R.; Ceriotti, M.; Chan, K.-T.; Cossio, P.; Dasetty, S.; Donadio, D.; Ensing, B.; Ferguson, A. L.; Fraux, G.; Gale, J. D.; Gervasio, F. L.; Giorgino, T.; Herringer, N. S. M.; Hocky, G. M.; Hoff, S. E.; Invernizzi, M.; Languin-Cattoën, O.; Leone, V.; Limongelli, V.; Lopez-Acevedo, O.; Marinelli, F.; Febrer Martinez, P.; Masetti, M.; Mehdi, S.; Michaelides, A.; Murtada, M. H.; Parrinello, M.; Piaggi, P. M.; Pietropaolo, A.; Pietrucci, F.; Pipolo, S.; Pritchard, C.; Raiteri, P.; Raniolo, S.; Rapetti, D.; Rizzi, V.; Rydzewski, J.; Salvalaglio, M.; Schran, C.; Seal, A.; Shayesteh Zadeh, A.; Silva, T. F. D.; Spiwok, V.; Stirnemann, G.; Sucerquia, D.; Tiwary, P.; Valsson, O.; Vendruscolo, M.; Voth, G. A.; White, A. D.; Wu, J. PLUMED Tutorials: a collaborative, community-driven learning ecosystem. arXiv preprint arXiv:2412.03595 2024,

(40) Bussi, G.; Gervasio, F. L.; Laio, A.; Parrinello, M. Free-energy landscape for *β* hairpin folding from combined parallel tempering and metadynamics. Journal of the American Chemical Society 2006, 128, 13435–13441.

