## Supplementary material for "Destabilization of Helix III Initiates Early Serum Amyloid A Misfolding by Exposing Its Amyloidogenic Core": The Supporting Information is available free of charge at https://pubs.acs.org.

### Devising Collective Variables

(a)

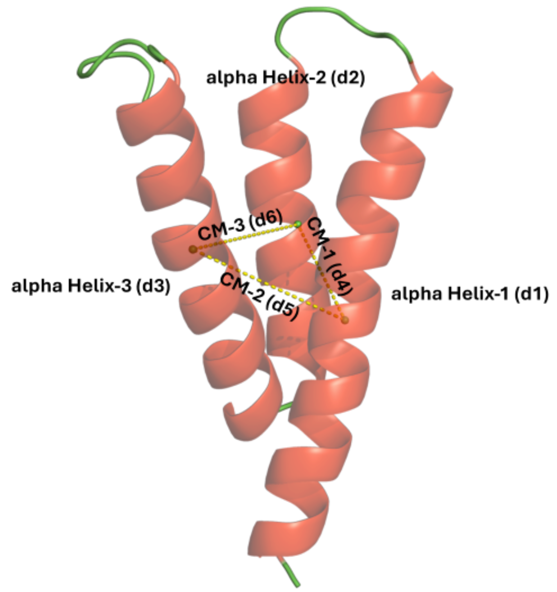


(b)

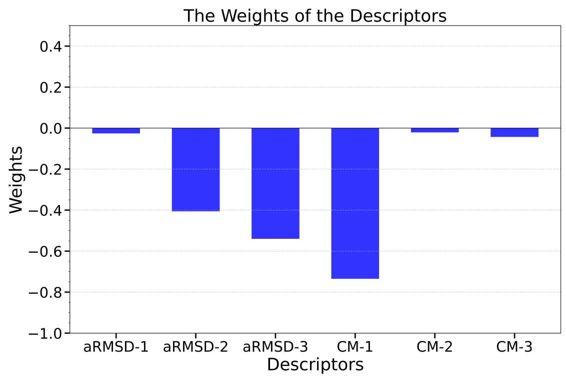


Figure S1: (a) Cartoon structure of $SAA_{1-76}$ with the three $\alpha$-helices shown in red; the distances between the centers of mass of the $\alpha$-helices used as descriptors are also indicated. (b) Weights assigned by HLDA to each of the descriptors.

### The Efficiency of the Simulation

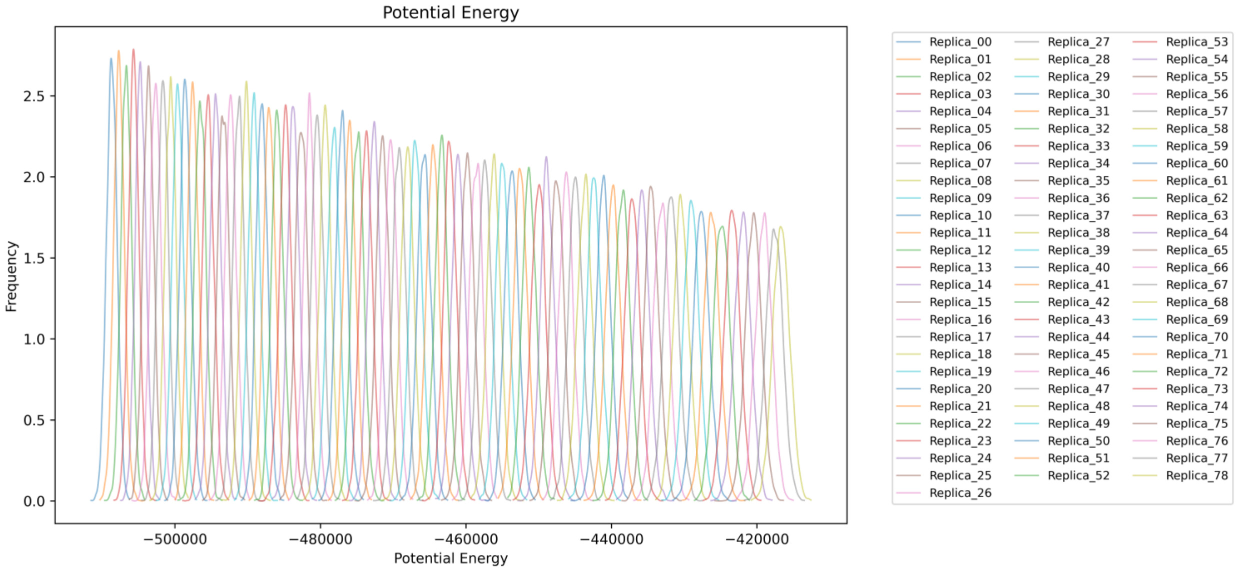


Figure S2: Potential energy probability distributions at all simulated temperatures

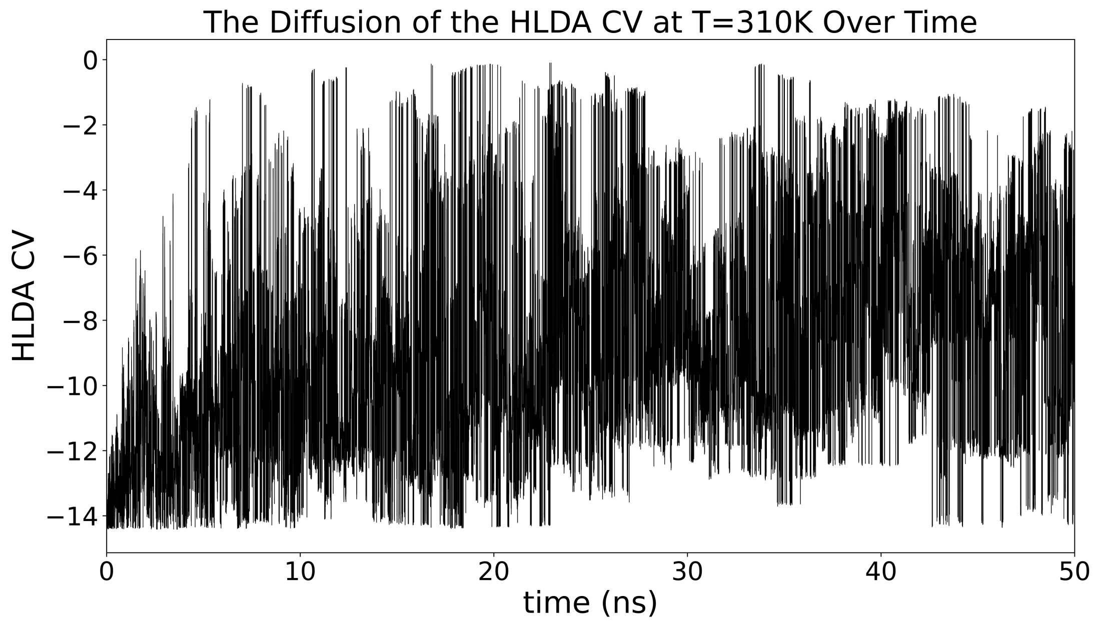


Figure S3: HLDA CV time dependence taken from T=310K replica of PTMetaD simulations.

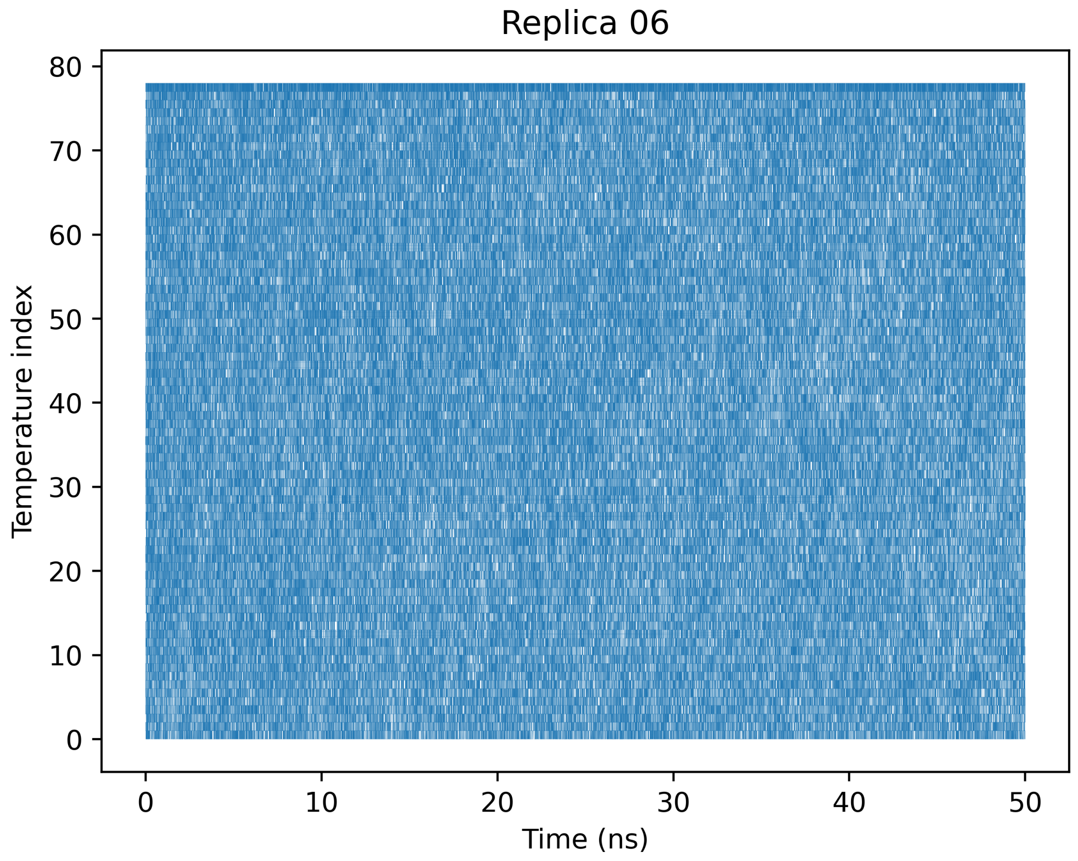


Figure S4: The time evolution of replica 6 through temperature space

#### The Convergence of the Simulation

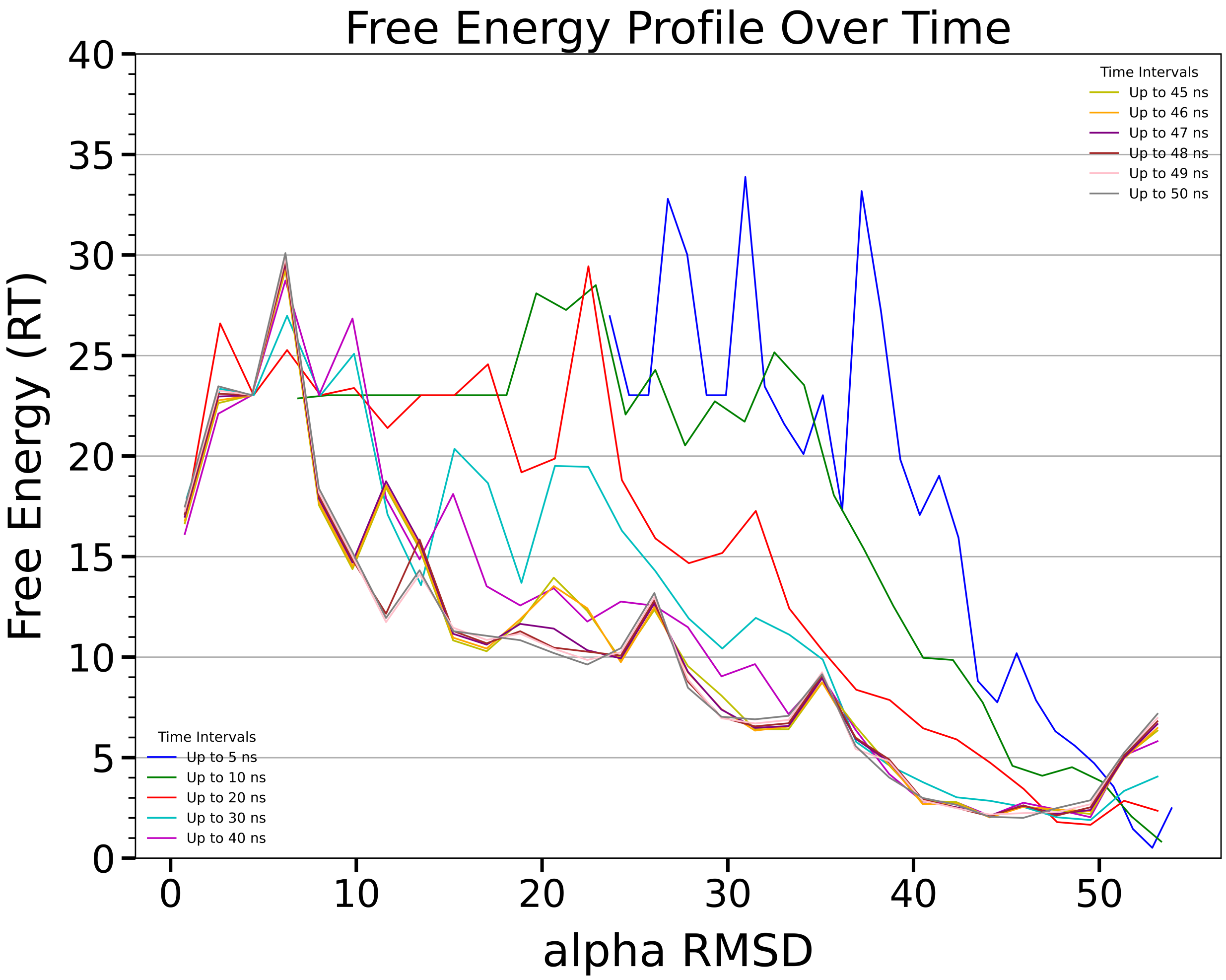


Figure S5: Overlapping of free energy profiles along $\alpha$RMSD CV in different time windows

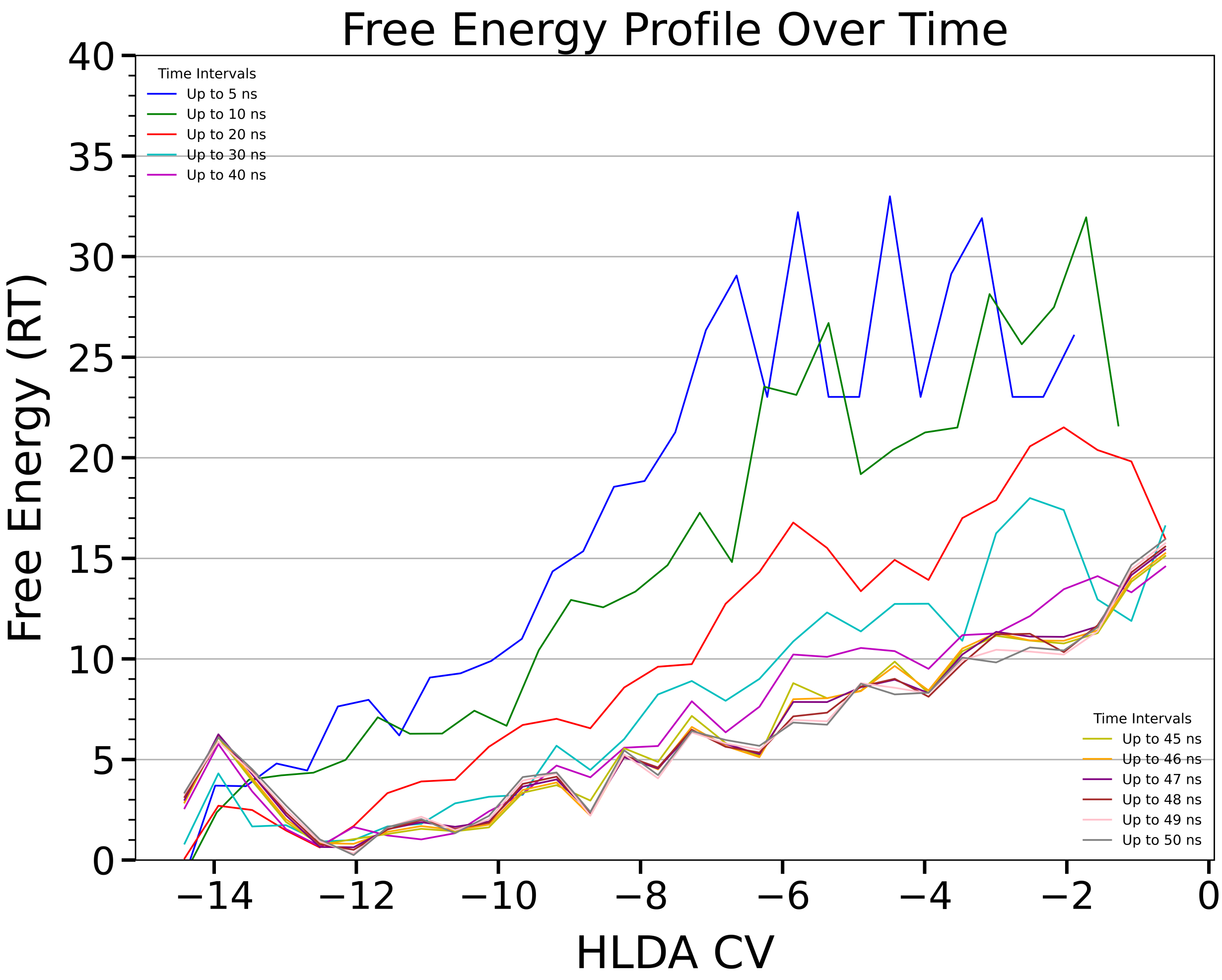


Figure S6: Overlapping of free energy profiles along HLDA CV in different time windows

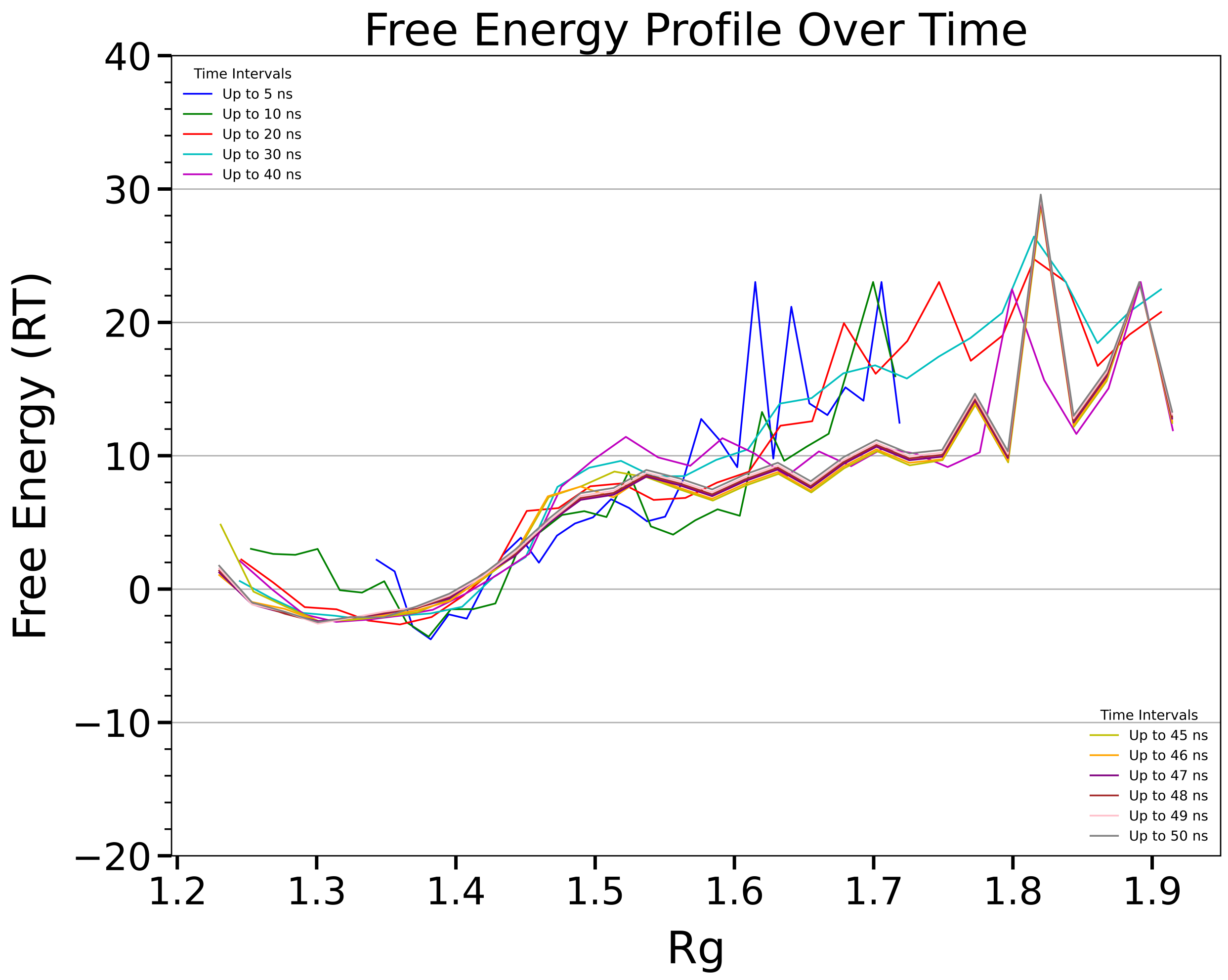


Figure S7: Overlapping of free energy profiles along Rg CV in different time windows

### SASA Analysis along the identified misfolding pathways

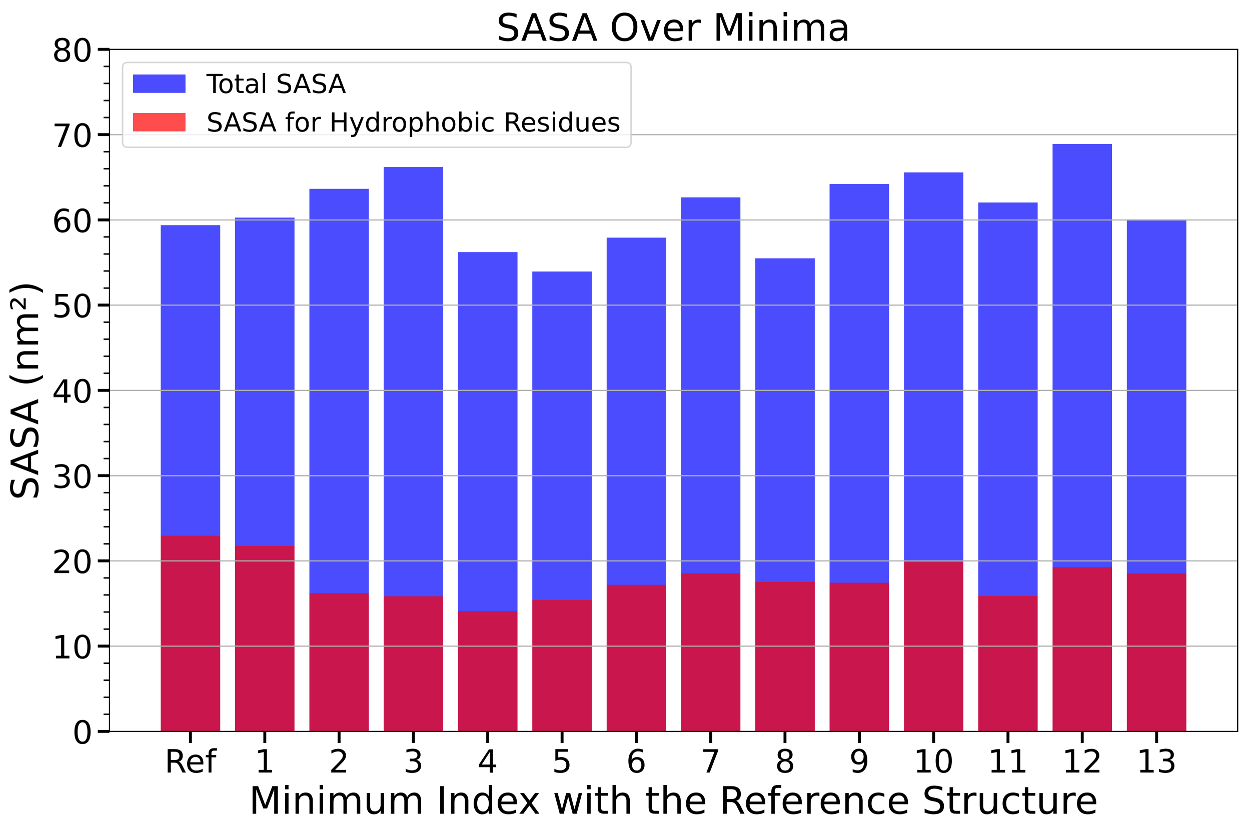


Figure S8: Total SASA for the protein and for the hydrophobic residues for the representative structure sampled during the simulations along aRMSD and HLDA-CV

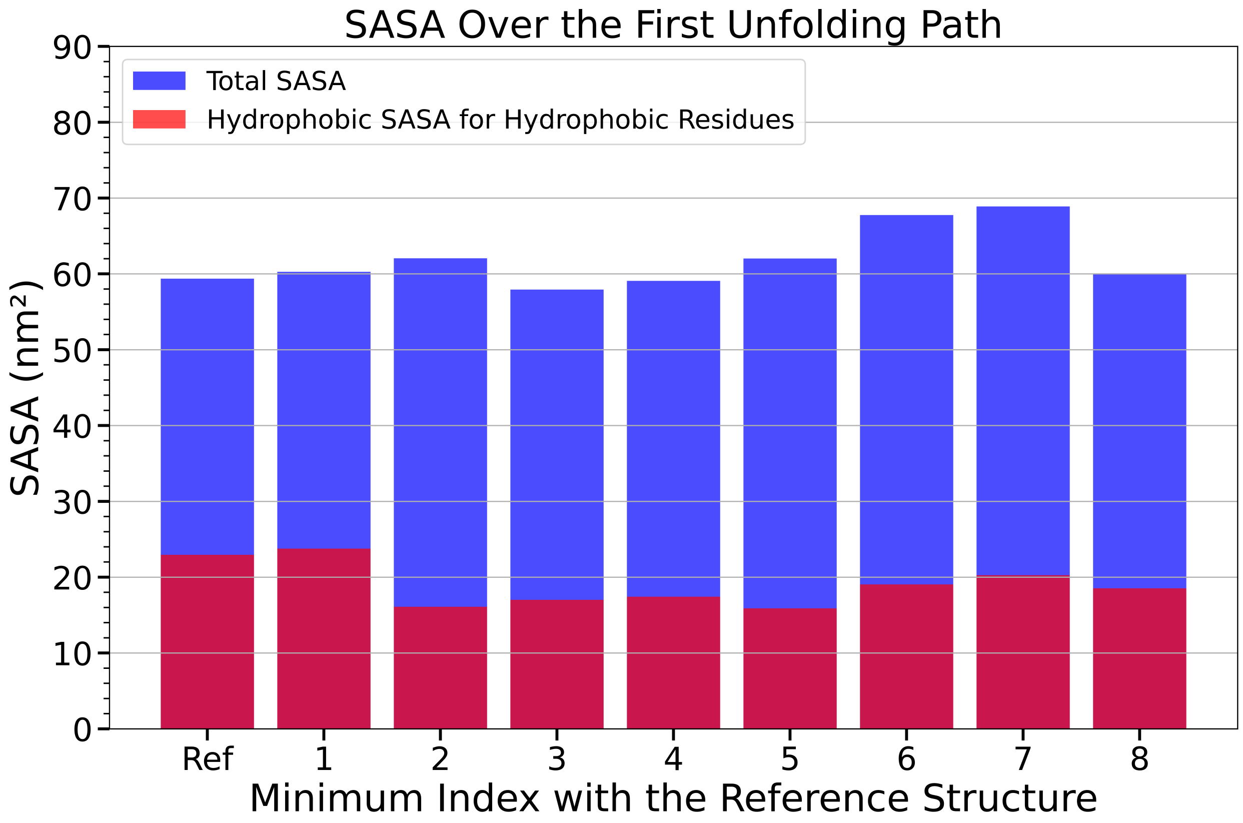


Figure S9: Total SASA for the protein and for the hydrophobic residues for the representative structure sampled during the simulations along the first misfolding path along aRMSD and Rg

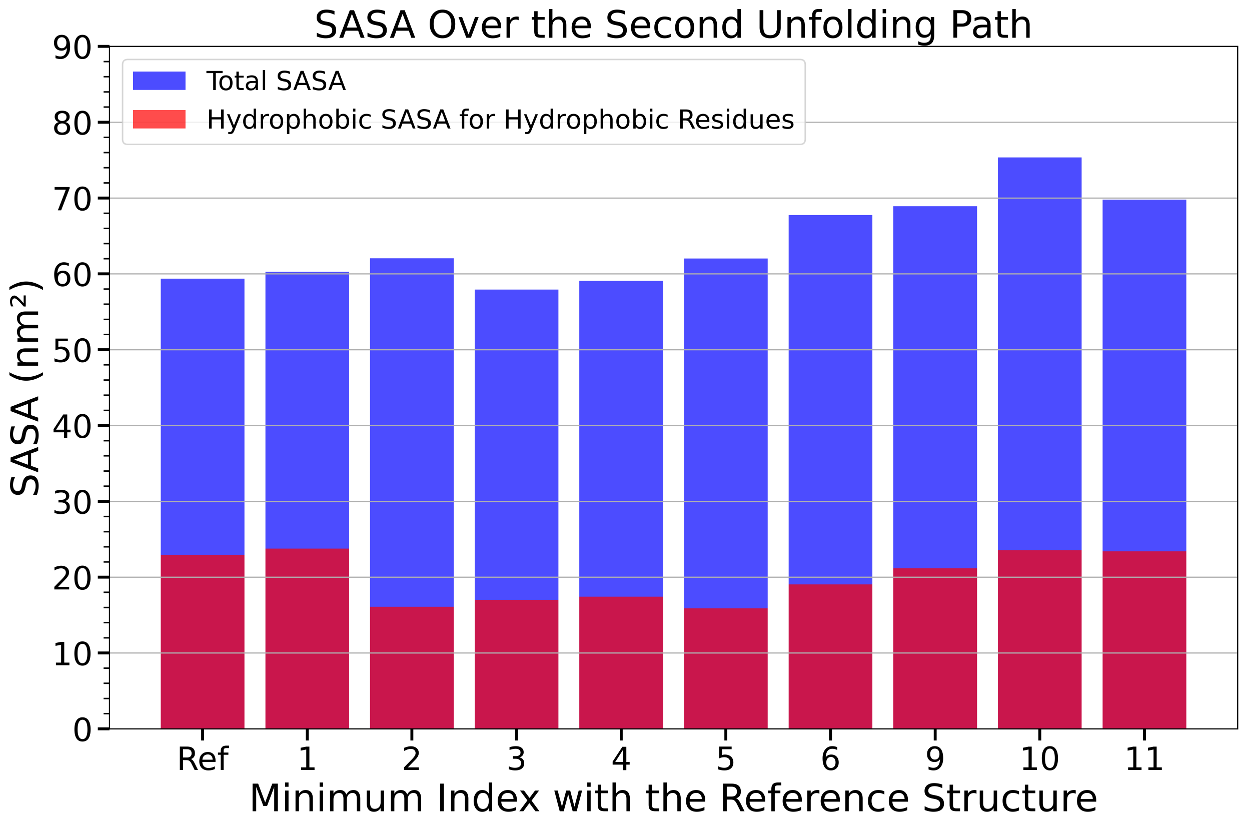


Figure S10: Total SASA for the protein and for the hydrophobic residues for the representative structure sampled during the simulations along the second misfolding path along aRMSD and Rg

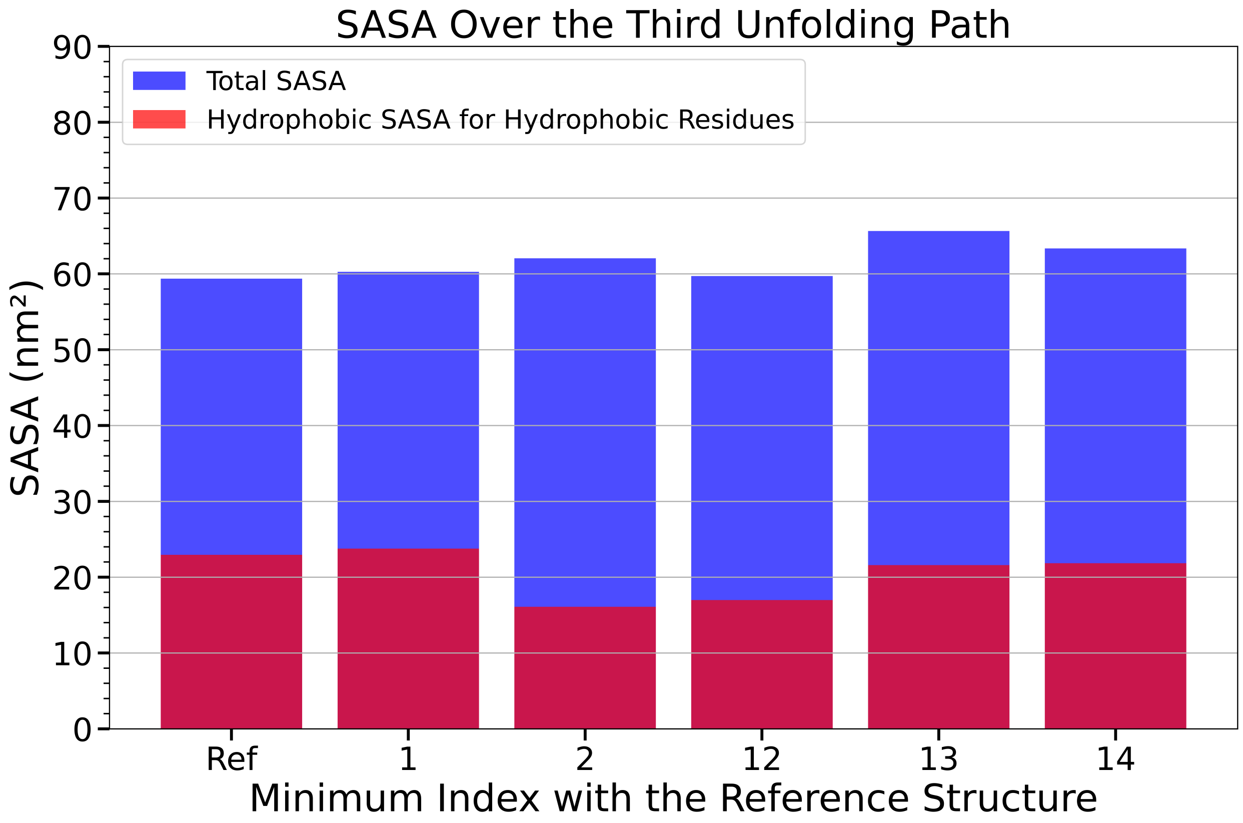


Figure S11: Total SASA for the protein and for the hydrophobic residues for the representative structure sampled during the simulations along the third misfolding path along aRMSD and Rg
